# Incorporation of unique molecular identifiers in TruSeq adapters improves the accuracy of quantitative sequencing

**DOI:** 10.1101/114603

**Authors:** Jungeui Hong, David Gresham

## Abstract

Quantitative analysis of next-generation sequencing data requires discriminating duplicate reads generated by PCR from identical molecules that are of unique origin. Typically, PCR duplicates are defined as sequence reads that align to the same genomic coordinates using reference-based alignment. However, identical molecules can be independently generated during library preparation. The false positive rate of coordinate-based deduplication has not been well characterized and may introduce unforeseen biases during analyses. We developed a cost-effective sequencing adapter design by modifying Illumina TruSeq adapters to incorporate a unique molecular identifier (UMI) while maintaining the capacity to undertake multiplexed sequencing. Incorporation of UMIs enables identification of bona fide PCR duplicates as identically mapped reads with identical UMIs. Using TruSeq adapters containing UMIs (TrUMIseq adapters), we find that accurate removal of PCR duplicates results in enhanced data quality for quantitative analysis of allele frequencies in heterogeneous populations and gene expression.

**Method Summary:** TrUMIseq adapters incorporate unique molecular identifiers in TruSeq adapters while maintaining the capacity to multiplex sequencing libraries using existing workflows. The use of UMIs increases the accuracy of quantitative sequencing assays, including RNAseq and allele frequency estimation, by enabling accurate detection of PCR duplicates.

---

Next-generation sequencing enables rapid and cost-effective identification of rare alleles from population of cells and quantitative expression profiling using RNA-seq. A critical technical issue in sequencing library preparation protocols, using either ligation-based or tagmentation-based approaches, is minimizing PCR duplicates that originate from library amplification prior to cluster generation (1-3). PCR duplicates represent redundant information that can inflate perceived read depth of specific genome or transcriptome sequences and therefore introduce biases in detecting minor frequency alleles in heterogeneous populations (4) or result in over-estimation of fragments derived from specific mRNAs.

In practice, PCR duplicates are removed using bioinformatics tools that detect duplicates based on the alignment information after mapping reads to a reference genome. However, one cannot know the rate of false positive or false negative duplicate detection using this method as there is no independent means of assessing whether an identical sequence read is the result of PCR amplification or reflects an independently generated molecule that is identical by chance.

Quantitative analysis of gene expression using RNA-seq, and the emergence of single cell mRNA sequencing, has emphasized the importance of identifying unique sequence reads for accurate quantitation of mRNA abundance using unique molecular identifiers (UMIs) to count unique molecules (5-10). A variety of methodological approaches to counting molecules have been developed for different sequencing applications. However, existing methods require specialized approaches and are not readily incorporated into existing workflows. For example, a design for amplicon sequencing that uses UMIs to detect PCR duplicates does so at the expense of removing the sample index and therefore multiplexing samples is no longer possible (11). A dual indexing adapter design, which includes both UMI and sample index, has been developed, but requires an additional sequencing phase resulting in additional sequencing cost (12). Alternatively, methodological advances have been introduced that minimize the impact of PCR duplicates. For example, optimizing PCR protocols can minimize PCR biases but requires extensive calibration (1, 3). PCR-free library preparation protocols avoid the problem of generating PCR duplicates, but remain limited in use due to the high cost of reagents and the requirement for greater amounts of starting material.

We developed a novel, cost-effective Illumina sequencing adapter design that enables identification of true PCR duplicates while maintaining the ability to perform library multiplexing. Our design incorporates both a sample index for multiplexed sequencing and a UMI for tagging unique molecules within a single sequencing adapter. Illumina TruSeq adapters are generated by annealing two partially complementary single-stranded oligonucleotides that typically contain a sample index for multiplexing (Figure 1A). In our design, we moved the multiplexing sample index to the 5’ end of the adapters proximate to the ligation site of the DNA insert and placed a 6 base pair UMI, by random incorporation of bases during oligonucleotide synthesis, at the position that typically contains the sample index. In principle, 4^6^, or 4096, UMIs are present in each TrUMIseq adapter. The method for preparing and sequencing libraries using TrUMIseq adapters is identical to methods for TruSeq adapters and thus readily implemented into existing workflows. Whereas different uniquely formed molecules may contain identical insert sequences or identical UMIs, the chance of both occurring is exceedingly rare and therefore, reads with identical mapping coordinates and identical UMI sequences are defined as true PCR duplicates (Figure 1B).

**Figure 1.**
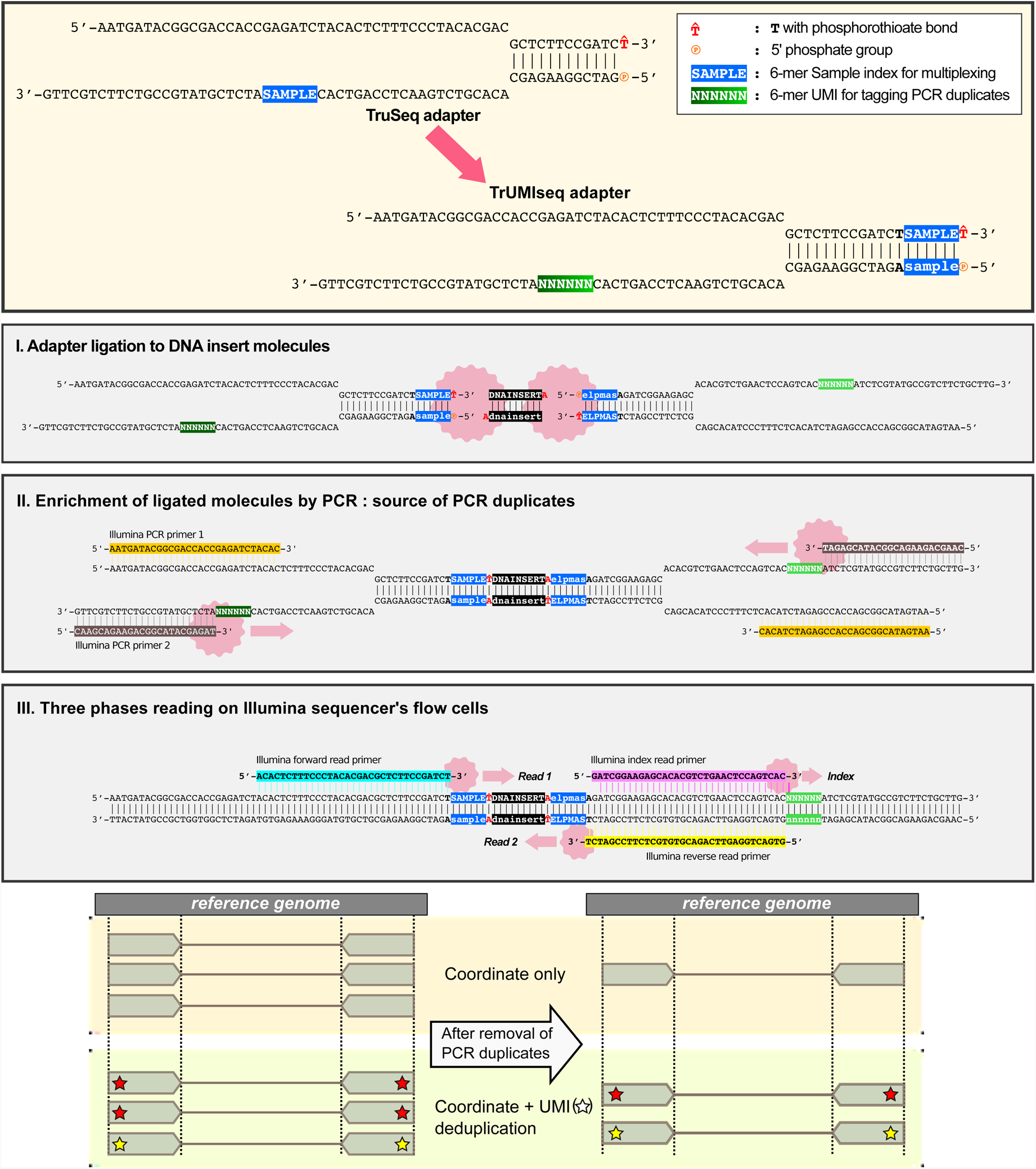
Accurate detection of PCR duplicates using TrUMIseq adapters. (**A**) TrUMIseq adapters are based on TruSeq adapters with relocation of the sample index and addition of a unique molecular identifier (UMI). Libraries are generated and sequenced with TrUMIseq adapters using the identical ligation, PCR and sequencing steps used for TruSeq adapters. (**B**). Removal of PCR duplicates using TrUMIseq adapters. Whereas coordinate-based deduplication depends on mapping information only, the use of UMIs enables distinction between true PCR duplicates that have identical UMIs (red star) from independently generated molecules that have different UMIs (yellow star).

We compared the rate of PCR duplicate detection using conventional genome coordinate-based detection with PCR duplicate detection using both mapping information and UMIs for three different sequencing protocols using TrUMIseq adapters: whole genome DNA sequencing (DNA-seq), targeted sequencing of amplicons (AMP-seq) and strand-specific RNA-seq using samples derived from *Saccharomyces cerevisiae.* The PCR duplicate rate is highly proportional to the number of PCR cycles used during library preparation and differs depending on the method of detection (Figure 2A). When considering only mapping information, the duplicate rate ranges from 20-40% for libraries prepared using less than 10 PCR cycles and up to 90% for libraries amplified with 15 cycles. By contrast, when the UMI information is incorporated to identify bona fide PCR duplicates, the duplication rate decreases to less than 10% for libraries constructed using less than 10 PCR cycles. Thus, up to 20% more unique sequencing reads can be recovered by using TrUMIseq adapters that would otherwise be incorrectly discarded without the use of UMIs.

**Figure 2.**
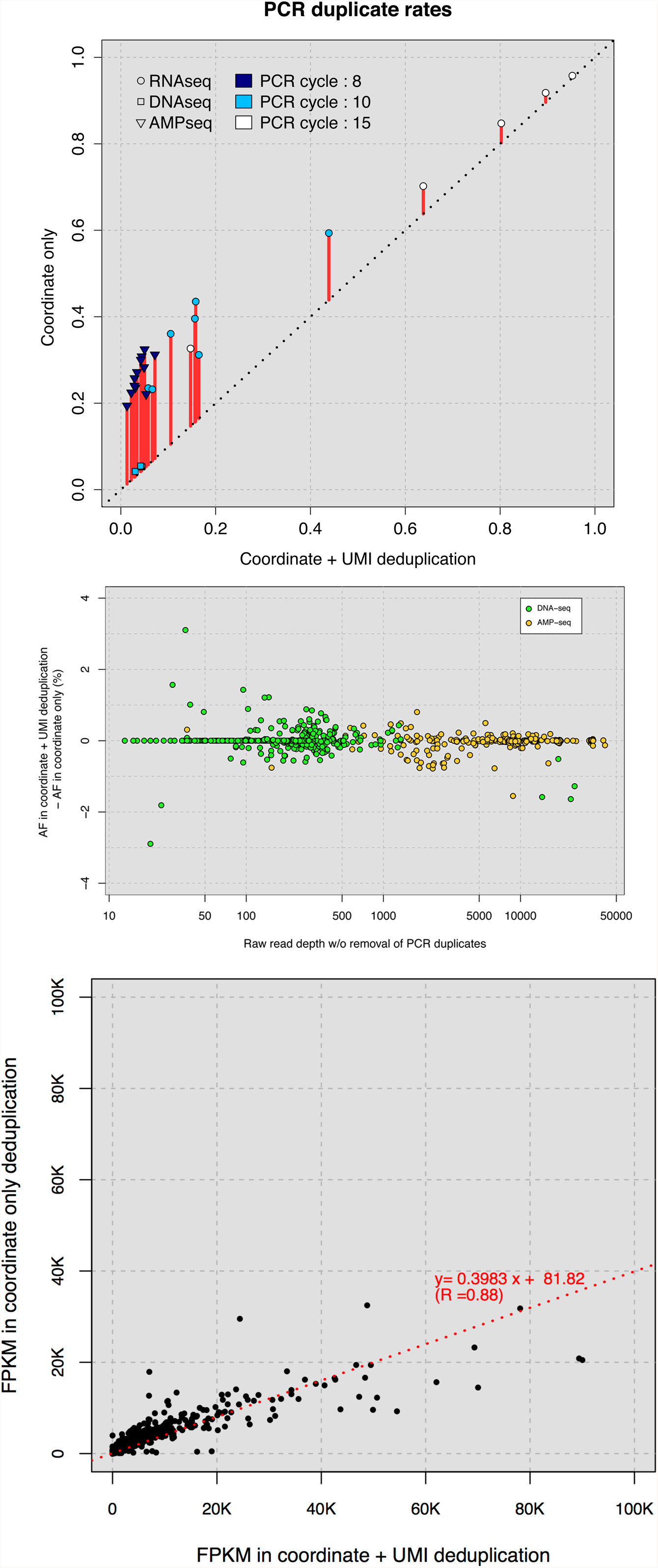
Accurate removal of PCR duplicates improves quantitative sequencing assays. (**A**) Comparison of PCR duplicates detection rates using mapping coordinates only and mapping coordinates with UMIs. (**B**) Differences in allele frequency estimates following deduplication using mapping coordinates only and using mapping coordinates in conjunction with UMIs. A total of 482 (DNA-seq) and 276 (Amp-seq) SNPs were studied. (**C**) Estimation of FPKMs for all yeast genes using mapping coordinate only deduplication compared with mapping coordinate and UMIs.

We find that each sequencing protocol differs in the estimated PCR duplicate rate. AMP-seq and RNA-seq data have very different estimated duplication rates using the two deduplication methods (triangle or circle data points in the Figure 2A). Interestingly, our DNA-seq data showed very low rate of PCR duplicates – less than 5% – regardless of the duplication detection method (rectangular data points in the Figure 2A). This is likely a combination of the fact that libraries from whole genomes are a more complex than libraries prepared from a subset of the genome (i.e. the transcriptome or targeted loci) and the large quantity of starting material used in our library preparation.

We investigated the effect of a reduced rate of false positive PCR duplicate detection using TrUMIseq adapters on quantifying rare allele frequencies in genome sequencing data. We compared differences in allele frequencies (AFs) of significant SNPs identified in DNA-seq and AMP-seq libraries for the multiple heterogeneous populations of *S. cerevisiae* following removal of PCR duplicates using UMI and coordinate information compared with the coordinate-only approach (Figure 2B). The majority of SNPs show less than 1% difference in their AF estimation using the two methods. However, the difference in estimated AF increases to up to 3% as read depth decreases in both types of samples suggesting that correctly identifying PCR duplicates is critical for identifying minor frequency alleles from heterogeneous population sequencing data.

We compared the impact of the two alternative approaches to RNAseq quantification of gene expression. Interestingly, we find that there is a clear effect of gene size on the differences in the resulting read count values: smaller genes tend to have greater loss of read counts when using coordinate-only deduplication (Supplementary Figure 1). This is likely due to the increased probability of generating an identical fragment for smaller genes. When considering all transcripts, the fragments per kilobase of transcript per million mapped reads (FPKM) values increase more than twofold when deduplication is performed using coordinate and UMI information (Figure 2C). Surprisingly, we find that estimates of differential gene expression for most genes are generally not affected by the use of coordinate only deduplication, although some down-regulated genes were sensitive to incorrect identification of PCR duplicates (Supplementary Figure 2). The bias in incorrectly identifying PCR duplicates in RNAseq data may be reproducible and thus comparison of fragment counts between samples may not be overly influenced by incorrect removal of duplicate reads. A non-random distribution of false positive PCR duplicates is supported by the good correlation (R = 0.88) in FPKM estimates between the two different methods of deduplication (Figure 2C).

One technical issue associated with TrUMIseq is that the presence of the sample index in the first part of read one results in low nucleotide diversity across a flow cell in the first 7 nucleotides of all reads. To address this issue, we suggest that multiple (at least four) libraries with diverse sample indices (Supplementary Table 1) should be pooled in a single sequencing lane. In addition, adding 5% or more PhiX spike-in is beneficial for increasing base complexity. Varying the length of sample indices would further increase base diversity.

Our results illustrate the utility of TrUMIseq adapters for distinguishing true PCR duplicates from randomly generated identical molecules. The procedure for library preparation and sequencing using TrUMIseq adapters uses existing TruSeq protocols, and primers, and therefore is readily implemented with, or alongside, existing workflows. TrUMIseq adapters are compatible with multiplexed sequencing in both paired end and single end sequencing modes requiring no additional sequencing reagents or primers unlike the dual indexing adapter design (12). TrUMIseq is also highly cost effective as it costs ~ $150 to make a stock of ~ 500μl of 20μΜ adapter, which can be used for constructing hundreds to thousands of libraries. The use of TrUMIseq adapters for accurate detection of PCR duplicates provides a readily implemented means of improving quantitative data quality for any sequencing application that currently uses TruSeq adapters.

## Author contributions

JH and DG conceived of the design. JH performed all experiments and computational analysis. JH and DG wrote the manuscript.

## Acknowledgements

We thank Tara Rock and the Genomic Core Facility at the New York University Center for Genomics and Systems Biology for assistance in implementing TrUMIseq and members of the Gresham lab for helpful discussions. This work was funded by the National Institute of Health (R01GM107466) and the National Science Foundation (MCB1244219). This paper is subject to the NIH Public Access Policy.

